# Characterization of alternative splicing in high-risk Wilms’ tumors

**DOI:** 10.1101/2023.10.30.564809

**Authors:** Yaron Trink, Achia Urbach, Benjamin Dekel, Peter Hohenstein, Jacob Goldberger, Tomer Kalisky

## Abstract

The significant heterogeneity of Wilms’ tumors between different patients is thought to arise from genetic and epigenetic distortions that occur during various stages of fetal kidney development in a way that is poorly understood. To address this, we characterized the heterogeneity of alternative mRNA splicing in Wilms’ tumors using a publicly available RNAseq dataset of high-risk Wilms’ tumors and normal kidney samples. Through Pareto task inference and cell deconvolution, we found that the tumors and normal kidney samples are organized according to progressive stages of kidney development within a triangle-shaped region in latent space, whose vertices, or “archetypes,” resemble the cap mesenchyme, the nephrogenic stroma, and epithelial tubular structures of the fetal kidney. We identified a set of genes that are alternatively spliced between tumors located in different regions of latent space and found that many of these genes are associated with the Epithelial to Mesenchymal Transition (EMT) and muscle development. Using motif enrichment analysis, we identified putative splicing regulators, some of which are associated with kidney development. Our findings provide new insights into the etiology of Wilms’ tumors and suggest that specific splicing mechanisms in early stages of development may contribute to tumor development in different patients.

## INTRODUCTION

Wilms’ tumors, also referred to as nephroblastomas, are malignancies affecting the kidney that primarily occur in children below the age of five. The tumorigenesis of Wilms’ tumors is believed to be closely associated with fetal nephron development since they often contain “blastemal” cells that histologically resemble fetal nephron progenitors. Moreover, Wilms’ tumors often arise from or in the vicinity of “nephrogenic rests” - poorly differentiated structures that resemble early embryonic kidney precursor cells not usually seen in the postnatal kidney [1–3]. These observations suggest that genetic and epigenetic aberrations during various stages of fetal kidney development lead to Wilms’ tumor formation, and that these aberrations result in extensive heterogeneity among patients.

There are two main protocols for treating Wilms’ tumors. The Children’s Oncology Group (COG) in North America recommends initial surgery followed by chemotherapy, whereas the International Society of Pediatric Oncology (SIOP), followed by European countries, favors initial chemotherapy prior to surgery [4,5]. Despite considerable improvement in treatment efficacy, patients with bilateral, relapsed, or “high-risk” tumors (see definition below) still present considerable challenges.

Wilms’ tumors are histologically and clinically diverse. Most cases are classified as Favorable Histology Wilms’ Tumors (FHWT), which are typically comprised of “stromal”, “blastemal”, and “epithelial” components. The *stromal* component resembles the nephrogenic stroma of the developing fetal kidney and may also contain cells resembling smooth or skeletal muscle, fat, cartilage, bone, and glial cells. The *blastemal* component histologically resembles the cap mesenchyme, a transient cell compartment of the fetal developing kidney which contains nephron progenitors [6]. In the fetal kidney, these nephron progenitors undergo a Mesenchymal to Epithelial Transition (MET) and differentiate into the various epithelial tubular segments of the nephron [7]. The *epithelial* component typically contains early non-functional epithelial structures from early stages of nephron differentiation. Tumors containing all three components and are classified as “mixed” type or “triphasic”, whereas those containing a single dominant histology are classified as “blastemal”, “stromal”, or “epithelial” tumors. The blastemal component is presumed to be most malignant, and tumors containing a large blastemal component that survived preoperative chemotherapy are regarded as “high-risk” tumors and require more aggressive treatment [6,8,9]. *Anaplastic* Wilms’ tumors are characterized by multipolar mitoses, nuclear enlargement, and hyperchromatic nuclei [10], and are classified as “focal” (FAWT) or “diffuse” (DAWT) depending on the geographical distribution of the anaplastic cells within the tumor [8,11]. Diffuse anaplastic Wilms’ tumors are associated with lower event-free survival and lower overall survival estimates when compared to Favorable Histology Wilms’ Tumors, and are also regarded as “high-risk” tumors [8,9,11]. Another class of high-risk tumors are favorable histology Wilms’ tumors that relapsed [8,12].

Previous analyses of Wilms’ tumors based on DNA microarrays or high throughput sequencing methods have been used to identify subsets of tumors based on gene expression patterns [13,14], to find differentially expressed genes between Wilms’ tumors and normal kidney samples [15,16], and to search for genetic markers associated with relapse [17,18]. In a previous study [19], we observed that Favorable Histology Wilms’ Tumors (treated according to the COG protocol) form a triangular shaped continuum in gene expression latent space, and that vertices of this triangle, which represent tumor “archetypes,” have blastemal, stromal, and epithelial characteristics, corresponding to the three main lineages of the developing fetal kidney. In a consecutive study [20] we found that this geometry is conserved also in high-risk tumors that were treated with chemotherapy prior to surgery (according to the SIOP protocol) but still contained a significant amount of remaining viable blastema [14], and used a probabilistic generative model [21] to represent each tumor as a mixture of three latent biological “topics” with stromal, epithelial, and blastemal characteristics.

While these studies focused on gene expression measurements, there is evidence that alternative mRNA splicing can also contribute to tumor initiation and progression. Recent studies identified sets of genes that are alternatively spliced in cancer [22–25], some of which are associated with the Epithelial to Mesenchymal Transition (EMT) [26,27] as well as poor clinical outcome [28]. Likewise, RNA binding proteins known to regulate mRNA splicing were found to be abnormally expressed or somatically mutated in cancer [29–33]. Specifically in Wilms’ tumor, a recent study showed that the splicing regulator ESRP2 is repressed by DNA methylation, whereas over-expression of ESRP2 in Wilms’ tumor cell lines promotes alternative splicing and inhibits cell proliferation both in-vitro and in-vivo [34]. In a recent work, we also found that fetal kidney cells at an early developmental stage have a mesenchymal splice-isoform profile that is similar to that of blastemal-predominant Wilms’ tumor xenografts [35]. Therefore, analysis of alternative splicing in different subtypes of Wilms’ tumors could provide insights to the molecular processes driving tumor development and their relation to kidney development.

In this study we set out to comprehensively characterize alternative splicing in high-risk Wilms’ tumors. We first downloaded a publically available dataset consisting of RNA sequences collected from favorable histology Wilms’ tumors (FHWT) that relapsed, diffuse anaplastic Wilms’ tumors (DAWT), and associated normal kidney samples [12]. These tumors were treated according to the COG protocol and collected by the NCI TARGET (“Therapeutically Applicable Research to Generate Effective Treatments”) initiative. Using Pareto task inference [36], Gene Ontology enrichment analysis, and cell deconvolution [37], we found that the tumors and normal kidney samples are organized according to progressive stages of kidney development within a triangle-shaped region in latent space, whose vertices, or “archetypes,” correspond to cell states resembling the cap mesenchyme, the nephrogenic stroma, and epithelial tubular structures of the fetal kidney. We next identified genes that are alternatively spliced between samples located near the three archetypes and found that many of these genes are related to EMT and muscle structure and development. Finally, we used motif enrichment analysis for known RNA binding proteins to identify putative splicing regulators, some of which were previously found in kidney development. We anticipate that these findings will contribute to a better understanding of the role of alternative mRNA splicing in the formation of Wilms’ tumors in diverse patient populations.

## RESULTS

### High-risk Wilms’ tumors and normal kidney samples form a triangle-shaped continuum in latent space that is bounded by archetypes with blastemal, stromal, and epithelial characteristics

We first downloaded an RNA sequencing dataset from the NCI TARGET [12] study containing 130 Wilms’ tumor samples and six associated normal kidney samples. The tumors were all classified as “high-risk” since they were either relapsed favorable histology Wilms’ tumors (FHWT) or diffuse anaplastic Wilms’ tumors (DAWT). We then performed sequence alignment and obtained a gene expression matrix (Table S1). Using Principal Components Analysis (PCA) we found that high-risk Wilms’ tumors form a continuum in gene expression latent space rather than discrete well-separated clusters [19,20] (Figure 1A-F). To better understand this continuous heterogeneity, we used Pareto Task Inference [36,38] to calculate the vertices of the best fitting polytope encompassing all tumors and normal kidney samples in latent space. These vertices, or “archetypes”, presumably represent idealized cell types from which all samples within the continuum are composed, where the precise transcriptional state of each sample determines its position in latent space relative to each biological archetype [19,36,39]. We tested different numbers of archetypes and found that three archetypes in two dimensions best describe the biological heterogeneity, whereas including an additional fourth archetype in the 3rd dimension captures variability that is likely related to RNAseq library size (see supplementary information).

**Figure 1:**
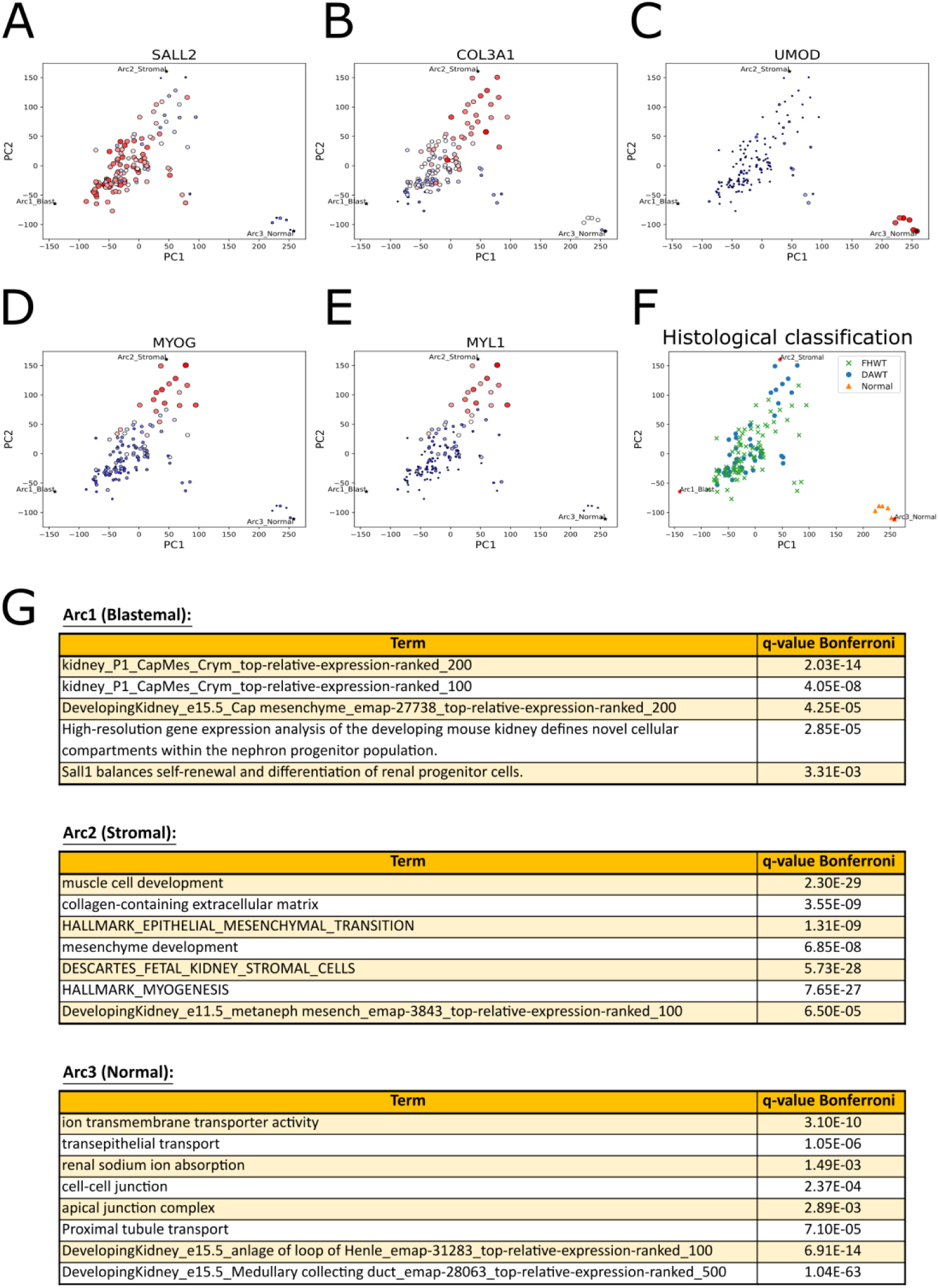
High-risk Wilms’ tumors and normal kidney samples form a triangle-shaped continuum in latent space that is bounded by archetypes with stromal, blastemal, and epithelial characteristics that resemble the nephrogenic stroma, the cap mesenchyme, and early epithelial structures of the fetal kidney, respectively. (A-C) Shown are PCA plots of RNAseq gene expression profiles from 130 high risk Wilms’ tumors (relapsed favorable histology or diffuse anaplastic Wilms’ tumors) and six normal kidneys from the US NCI TARGET project. Three archetypes, labeled “Arc1”, “Arc2”, and “Arc3”, form the vertices of the best fitting polytope which encompasses all data points. To identify the archetypes, the size and color of each sample (dot) are drawn according to expression levels of known genes marking cell population of the fetal developing kidney (large red – high expression, small blue – low expression). The gene SALL2, which marks the cap mesenchyme, is highly expressed near Arc1 which is the “blastemal” archetype. The gene COL3A1, which marks the nephrogenic stroma, is highly expressed near Arc2 which is the “stromal” archetype. Likewise, UMOD a gene which is highly expressed in the epithelial cells of the loop of Henle, is highly expressed in the normal samples located near Arc3, which is the “normal” archetype that is predominantly epithelial. (D,E) MYOG, a transcription factor that can induce myogenesis, and MYL1, a gene that encodes a muscle motor protein, are highly expressed near the “stromal” archetype (Arc2). (F) A PCA plot with each sample marked according to its histological classification. We did not observe a significant separation between relapsed favorable histology Wilms’ tumors (FHWT) and diffuse anaplastic Wilms’ tumors (DAWT) in this dataset. (G) Genes overexpressed (log2FC > 3) in each of the three archetypes with respect to the other two were used as input for gene ontology enrichment analysis. Arc1, the “blastemal” archetype, overexpresses genes related to the cap mesenchyme and nephron progenitor cells. Arc2, the “stromal” archetype, overexpresses genes that are related to the extracellular matrix, the nephrogenic stroma (the un-induced metanephric mesenchyme), and muscle development. Arc3, the “normal” archetype, overexpresses genes that are related to the structure and function of epithelial lineages of the kidney.

To infer the biological identity of the three archetypes, we checked the expression levels of selected genes that are known to mark the different lineages in the developing kidney (Figures 1A-C, Figure 2A). We found that the gene SALL2, a marker for the cap mesenchyme, is over-expressed near the first archetype, hence we named it as the “blastemal” archetype. Similarly, the gene COL3A1, which is a marker of the nephrogenic stroma, is overexpressed in the vicinity of the second archetype, which we named the “stromal” archetype, and the gene UMOD, which is a marker for renal epithelial cells, is over-expressed in the normal samples near the third archetype, which we named the “normal” archetype. We also observed that MYOG, a muscle-specific transcription factor with myogenesis-inducing capabilities, and MYL1, a motor protein expressed in muscle cells, are overexpressed near the stromal archetype (Figures 1D-E). Consistent with the findings of the original paper by the TARGET initiative [12], we did not observe an overall significant separation in gene expression latent space between tumors with favorable histology (FHWT) and diffuse anaplasia (DAWT) (Figure 1F).

**Figure 2:**
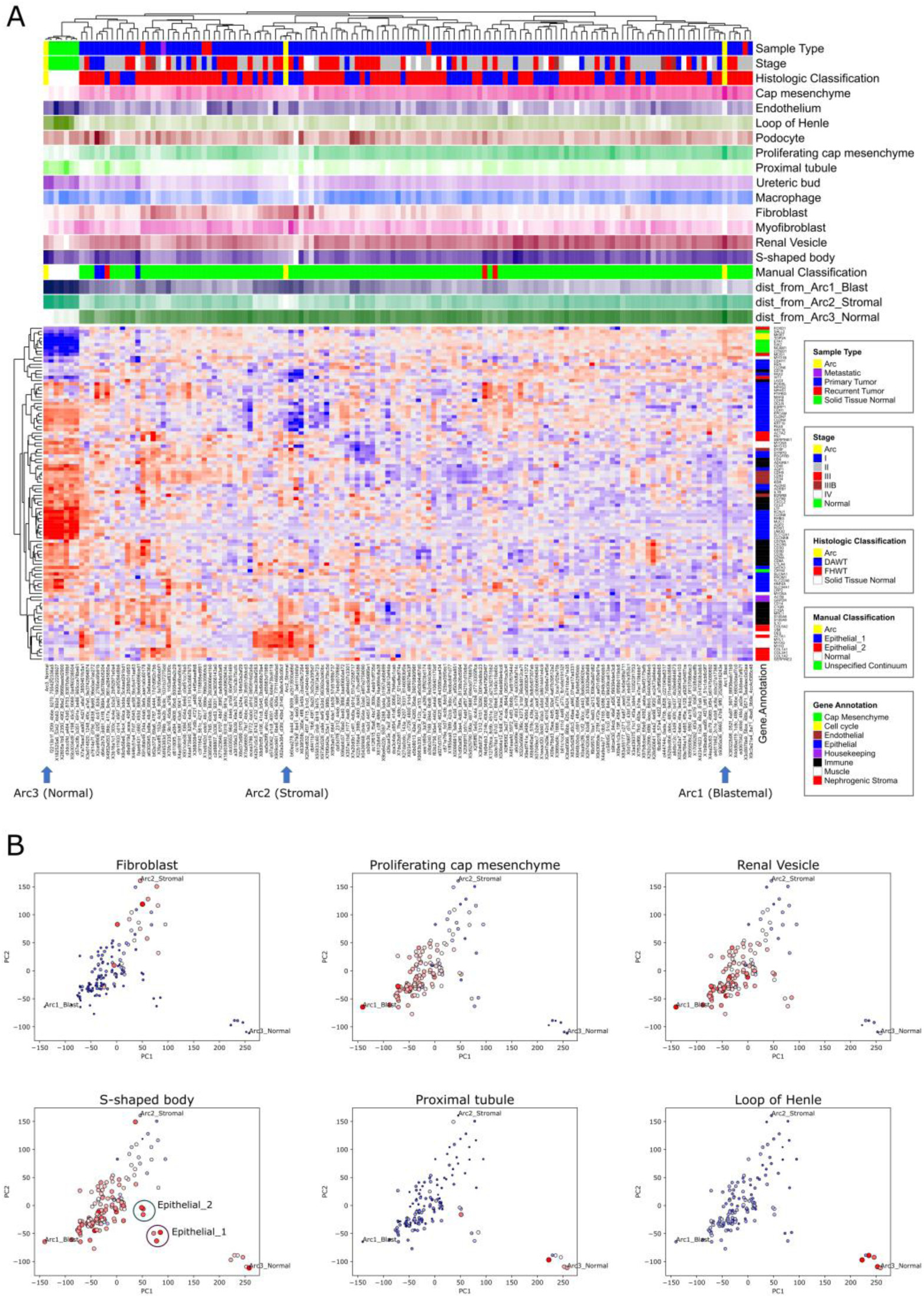
Cell deconvolution shows that high risk Wilms’ tumors are organized in latent space according to progressive stages of kidney development. (A) A gene expression heatmap of 102 selected genes that are known from the literature to be involved in kidney development. The top panels show the clinical and histological information for each sample, the proportions of cell types within each sample as predicted by cell deconvolution, and the distances from each of the three archetypes. It can be seen that genes characteristic to the cap mesenchyme (e.g. CITED1, EYA1, and SALL2) are overexpressed near the blastemal archetype. Genes characteristic of the nephrogenic stroma (e.g. COL3A1, COL5A2, and VIM), as well as muscle structure and development (e.g. DES, MYL1, and MYOG), are overexpressed in the vicinity of the stromal archetype. Likewise, genes marking epithelial tubular structures, mainly the loop of Henle and the distal tubule (e.g. CDH1, KCNJ1, MUC1, SLC12A1, and UMOD), are overexpressed near the normal archetype. (B) PCA plots marking proportions of fetal kidney cell populations within each sample, as inferred by cell deconvolution. The size and color of each data point are drawn according to the proportion of the selected population from the predicted cellular composition (large red – high proportion, small blue – low proportion). It can be seen that the samples are arranged in latent space according progressive stages of the developing kidney: cells resembling the cap mesenchyme are predominant in tumors located near the blastemal archetype, whereas cells resembling early nephron differentiation such as the “renal vesicle” and “S-shaped body” are found at progressively higher proportions in the more epithelial-like tumors that are located towards the normal archetype (“Epithelial 1” and “Epithelial 2”). Similarly, cells resembling fully differentiated epithelial structures such as the proximal tubule and the loop of Henle are predominant in the normal samples. Likewise, we observed that tumors located near the stromal archetype have a high proportion of cells resembling renal fibroblasts, which are the predominant component of the nephrogenic stroma.

We further characterized the identity of the archetypes by performing gene ontology (GO) enrichment analysis (Figure 1G, Table S2). We chose genes that are over-expressed (log2FC > 3) in each of the three archetypes with respect to the other two and used these as inputs to ToppGene [40]. We observed that genes that are overexpressed in the blastemal archetype are related to the cap mesenchyme and nephron progenitors, genes that are overexpressed in the stromal archetype are related to the nephrogenic stroma (the un-induced mesenchyme) and muscle development, and genes that are enriched in the normal archetype are related to more differentiated epithelial structures of the nephron such as the proximal tubule, the loop of Henle (LOH), and the collecting duct.

### High risk Wilms’ tumors are organized in latent space according to progressive stages of kidney development

In order to better understand the relationship between Wilms’ tumor heterogeneity and kidney development, we performed hierarchical clustering using expression levels of 102 selected genes that are known from the literature to be involved in kidney development (Figure 2A, Table S3, supplementary information). We observed that indeed, the tumors near the stromal archetype have high expression of genes related to the nephrogenic stroma (e.g. COL3A1, COL5A2, and VIM) and muscle structure and development (e.g. DES, MYL1, and MYOG), tumors located near the blastemal archetype have high expression of genes marking the cap mesenchyme (e.g. CITED1, EYA1, and SALL2). Likewise, we observed that the normal samples have high expression of renal epithelial markers, mainly for the loop of Henle and the distal tubule (e.g. CDH1, KCNJ1, MUC1, SLC12A1, and UMOD). Finally, we identified six “epithelial-like” tumors that are located distinctly from the others in latent space (marked “Epithelial 1” and “Epithelial 2” in Figure 2B) and that also express some renal epithelial markers (e.g., CDH6, KRT18, and KRT19, see supplementary information).

We next used cell deconvolution [37] to predict the proportions of different cell types in each sample. Since Wilms’ tumors contain cells resembling those of the fetal kidney, we used a publicly available atlas of single cell gene expression from the human fetal kidney [41] as a reference. After deconvolving each sample, we plotted the proportions of each predicted fetal cell subpopulation (e.g. “fibroblast”, “proximal tubule” etc.) for each sample in gene expression latent space (Figures 2B and supplementary information). We observed that the tumors are arranged according to progressive developmental stages in latent space. For example, high proportions of cells resembling the cap mesenchyme are found within tumors located near the blastemal archetype, whereas cells resembling early stages of nephron differentiation (“renal vesicle” and “S-shaped body”) are found in progressively higher proportions in the more epithelial-like tumors that are located towards the normal archetype (marked as “Epithelial 1” and “Epithelial 2” in Figure 2B). Cells resembling fully differentiated epithelial structures (“proximal tubule” and “loop of Henle”) were found to be predominant in the normal samples. Likewise, we observed that tumors located near the stromal archetype have a high proportion of cells resembling renal fibroblasts. We performed similar analysis on a dataset of high-risk blastemal-type Wilms’ tumors (treated according to SIOP protocols) published by Wegert et al. [14] and found similar results [20] (see a detailed comparison in the supplementary information).

### A significant number of RNA transcripts related to EMT and muscle development are alternatively spliced between high risk Wilms’ tumors located in different regions of latent space

We next set to identify genes that are alternatively spliced between the different tumors and normal kidney samples located in different regions in latent space. We first chose three groups of samples, each containing five samples located near one of the three archetypes (see supplementary information). We then used rMATS [42] to perform three comparisons between these three groups and found a set of transcripts that are significantly alternatively spliced (Figures 3A-B, Tables S4-S5, and supplementary information). Gene Ontology (GO) enrichment analysis for alternatively skipped exons (FDR = 0, |Δ*ψ*| > 0.1), showed that a significant fraction of these transcripts are related to the Epithelial to Mesenchymal Transition (EMT) and muscle development (Figure 4A, Table S4).

**Figure 3:**
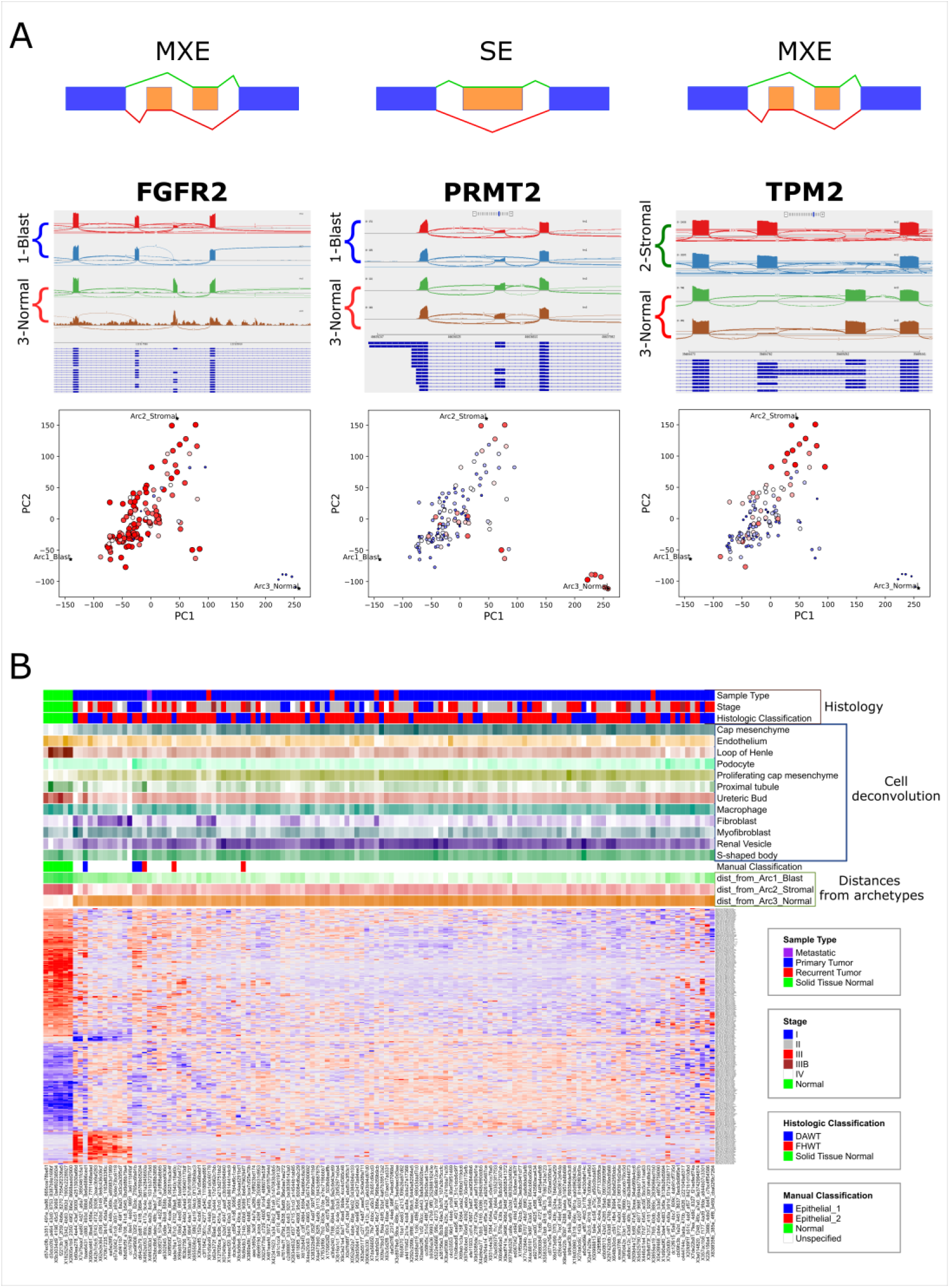
RNA transcripts related to EMT and muscle development are alternatively spliced between high risk Wilms’ tumors located in different regions of latent space. (A) Sashimi plots (top) and feature plots (bottom) for the alternatively spliced genes FGFR2, PRMT2, and TPM2. The sashimi plots show representative samples that are located near the archetypes. The feature plots show the isoform inclusion levels in the different tumors and samples in latent space, where the size and color of each sample are drawn according to the inclusion levels of a particular isoform (large red – high, small blue - low). (B) A heatmap showing the inclusion levels of 277 significantly skipped/included exons (SE) and mutually exclusive exons (MXE) for transcripts that were found to be significantly alternatively spliced in latent space. The rows are the union of all significant alternative splicing events (FDR = 0, |Δ*ψ*| > 0.2) from each of the three comparisons between the five nearest neighbors of the three archetypes. We plotted only SE (single-exon) and MXE (mutually exclusive exons) splicing events since these were the majority. The top panels show clinical and histological information, the proportions of cell types in each sample as predicted by cell deconvolution, our manual classification based on tumor location in latent space, and the distances to each of the three archetypes.

**Figure 4:**
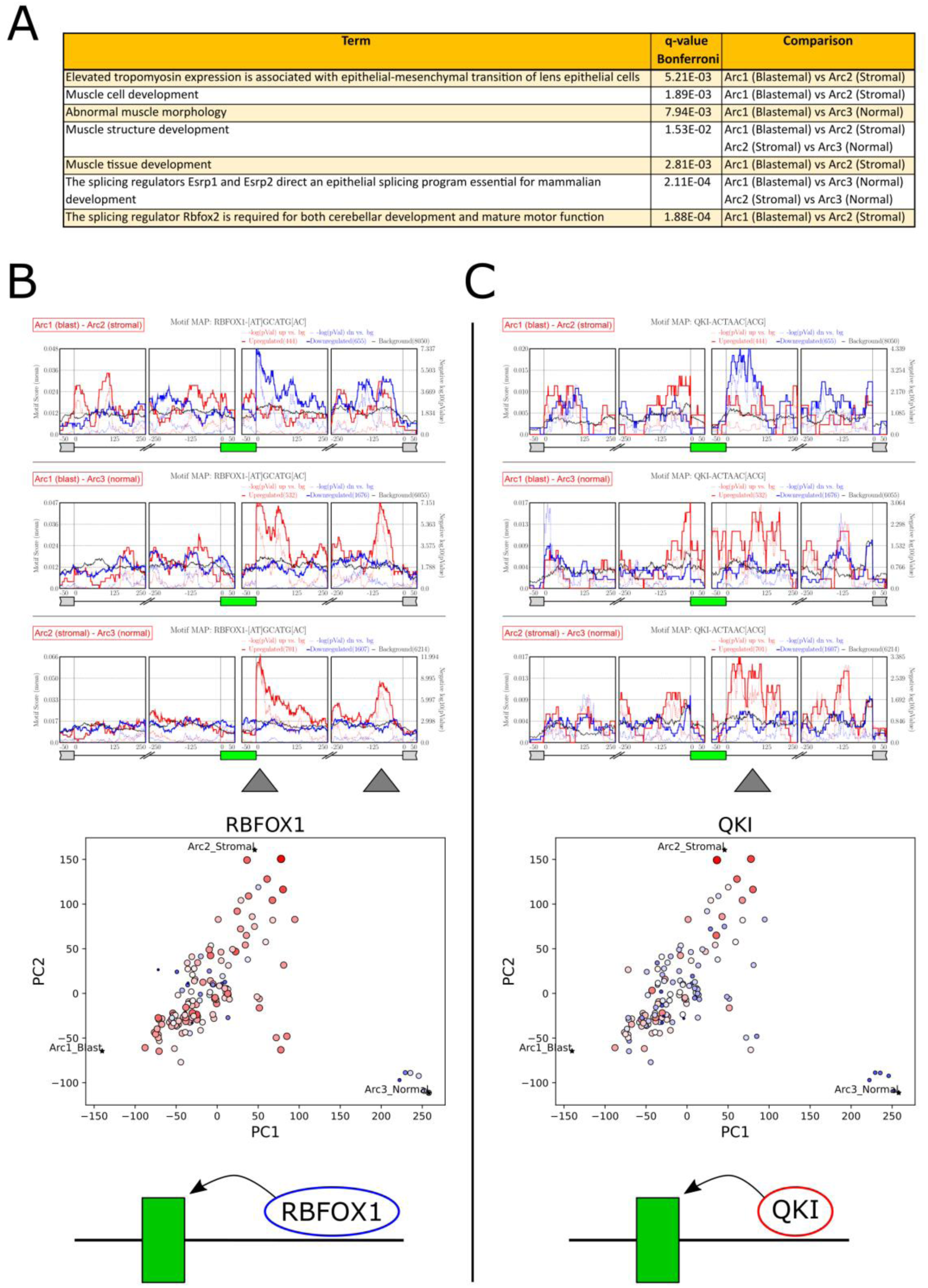
Motif enrichment analysis of RNA binding proteins reveals putative RNA binding proteins (RBPs) regulating alternative splicing between Wilms’ tumors located in different regions of latent space. (A) Gene Ontology (GO) enrichment analysis for exons that were found to be significantly alternatively spliced (FDR = 0, |Δ*ψ*| > 0.1) in latent space, as identified from the three comparisons between the five nearest neighbors of the three archetypes. A significant fraction of these transcripts are related to the Epithelial to Mesenchymal Transition (EMT) and muscle development. (B, C) Motif enrichment diagrams (top) and expression feature plots (middle) for the splicing regulators RBFOX1 and QKI. It can be seen that the expression levels these RNA binding proteins are elevated in the blastemal, and even more in the stromal tumors. Independently, we observed that exons that were elevated in these tumors were enriched for binding motifs of these regulators at their downstream introns. This indicates that in these tumors, RBFOX1 and QKI bind to the mRNA downstream of the mesenchymal associated cassette exons and promote their inclusion (bottom sketch).

For example, the gene FGFR2, a gene that is known to have epithelial and mesenchymal splice variants [43–46], is expressed in its mesenchymal form in the majority of tumors. Similarly, the gene PRMT2, a gene that is involved in cancer invasion, growth, and presumably EMT, and whose splice variants have been linked to cancer [47,48], contains an exon that is over-expressed near the stromal and normal archetypes, as well as some epithelial-like tumors. Likewise, the gene TPM2, whose splice variants and differential expression have been associated with EMT [49,50], cancer [51,52], and muscle diseases [53], is expressed in its muscle-specific isoform [51,54] near the stromal archetype. Another example is the gene FLNB whose exon 30 was found to be skipped in the majority of tumors (see supplementary information), which is consistent with previous findings that skipping of this exon induces EMT, is associated with basal-like breast cancer, and is regulated by the RNA binding proteins QKI and RBFOX1 [55]. Similarly, in the gene ATP2B1 (PMCA1) an isoform that is over-expressed specifically in skeletal and heart muscle, but not in kidney tissues [56,57], was found to be over-expressed near the stromal archetype (see supplementary information). Another notable example is the gene MEF2D [58], whose muscle-specific isoform is over-expressed near the stromal archetype (see supplementary information). The muscle-specific isoform for this gene was previously found to be promoted by the splicing regulators RBFOX1/2 [59,60].

### Motif enrichment analysis of RNA binding proteins reveals putative RNA binding proteins (RBPs) regulating alternative splicing between Wilms’ tumors located in different regions of latent space

GO enrichment analysis for transcripts that are alternatively spliced between samples located near the three archetypes suggests that many of them are known targets for the splicing regulators RBFOX2, ESRP1, and ESRP2 (Figure 4A). Therefore, in order to identify more putative splicing regulators we used rMAPS [61,62] to perform enrichment analysis for RNA binding motifs belonging to known RNA binding proteins (Figure 4B-C, Table S6, and supplementary information). We found a set of putative splicing regulators whose binding motifs were enriched upstream or downstream of exons that were alternatively spliced, and that were also differentially expressed between the samples in different locations in latent space. The most prominent motif enrichment was found for the RNA binding proteins RBFOX1 [55,63] and QKI [55,63,64], which indicates that in the more stromal tumors, these splicing regulators are over-expressed and tend to bind downstream of their target exons and promote their inclusion. We note that RBFOX1 and QKI were previously shown to physically interact with each other in order to regulate alternative splicing in mammary epithelial cells, thereby promoting an epithelial to mesenchymal transition (EMT) [55], and have also been associated with the mesenchymal to epithelial transition (MET) that occurs during fetal kidney development [35].

## DISCUSSION

In this study we characterized the continuous heterogeneity of high-risk Wilms’ tumors using gene expression and mRNA splicing. The development of Wilms’ tumors has been tightly linked to aberrations in fetal kidney development. Two central processes in embryonic nephrogenesis are the induction of the metanephric mesenchyme, resulting in condensation of cells from the nephrogenic stroma around the ureteric tip to form the cap mesenchyme, and the mesenchymal to epithelial transition (MET), by which cells from the cap mesenchyme progressively differentiate through a series of transformations to form the various epithelial segments of the nephron. Here we first showed that high-risk Wilms’ tumors and normal kidney samples from different patients form a continuum in gene expression latent space that is bound by stromal, blastemal, and epithelial archetypes that resemble the nephrogenic stroma (the un-induced metanephric mesenchyme), the cap mesenchyme, and early epithelial structures of the fetal kidney, respectively. Then, we inspected alternative mRNA splicing between tumors along this continuum in latent space and found a set of transcripts that are alternatively spliced, many of which are related to EMT and muscle development. In particular, we observed elevated expression of muscle-specific isoforms in tumors located near the stromal archetype. Since both the kidney metanephric mesenchyme and muscle tissues originate from the fetal mesoderm, there are two possible hypotheses: either that the more stromal Wilms’ tumors were distorted in early stages of development at the mesoderm stage, before the onset of kidney development, or that after initial differentiation into an early kidney lineage they had undergone dedifferentiation or transdifferentiation to a muscle-like lineage. Future functional studies will be required to test these possibilities. We believe that our results have the potential to assist in pinpointing the developmental trajectory of Wilms tumors in patients and to facilitate the development of novel therapies aimed at targeting specific mRNA splicing mechanisms associated with this disease.

## METHODS

### Datasets and preprocessing

136 BAM files from the TARGET Wilms’ Tumor study [12] were downloaded from the Genomics Data Commons (GDC) data portal using the GDC Data Transfer Tool Client (access date: December 1, 2020). Each BAM file was sorted and converted to a paired-end fastq file using SAMtools [65] and realigned to a reference genome (hg38) using STAR [66] to produce a gene expression counts matrix. Clinical data was manually downloaded from the GDC website. The raw gene expression counts were normalized using DESeq2 [67] and then a modified log-transform was performed (log2(1+x)). Genes with zero counts in all samples were removed from the analysis.

### Data visualization, GO enrichment analysis, and clustering

PCA was performed using the “Scikit-learn” python package [68] and PCA plots were drawn using the Matplotlib python package [69]. Gene Ontology enrichment analysis was performed using ToppGene (Chen et al. 2007). Hierarchical clustering was done using the “ComplexHeatmap” R package [70], with standardized rows (=genes). For clustering of gene expression, we used the Pearson correlation distance metric with complete linkage. For clustering of mRNA splicing inclusion levels we used the Euclidean distance metric with average linkage.

### Archetype Analysis

The “ParTI_lite” MATLAB function [36] was used to find the best fitting polytope encompassing the data points in latent space, where each data point is the normalized gene expression vector of a specific sample. The vertices of the best fitting polytope represent archetypes - idealized tumors/tissues or cell types which specialize at a particular biological task. The “performance” of each sample at this task is determined by its geometrical distance from each of the archetypes, and the identity of the biological tasks that characterize each archetype can be inferred from genes enriched in nearby latent space [36].

### Cell deconvolution

For cell deconvolution we used the CPM [37] algorithm as implemented in the “scBio” R package. We used the human fetal kidney single-cell RNAseq dataset from the Kidney Cell Atlas [41] as a reference single cell gene expression matrix and a UMAP [71] embedding of this dataset as the cell state space. We used the default values for the “modelSize”, “minSelection”, and “neighborhoodSize” parameters, and set the parameter “quantifyTypes” = T, which instructs the algorithm to quantify the proportions of different reference cell sub-populations in each bulk sample, in addition to the abundance values of each reference single-cell.

### Alternative splicing and RNA binding motif enrichment analysis

We used rMATS [42] to find mRNA splicing events with significant inclusion level differences between groups of samples located near each of the archetypes. We manually inspected top alternatively spliced transcripts in the IGV genome browser [72]. We used the tables output by rMATS as input to rMAPS [61,62] in order to identify putative splicing regulators by searching for binding motifs belonging to known RNA binding proteins (RBPs) that are also enriched in the vicinity of alternatively spliced exons. Additionally, in order to draw the heatmap of inclusion levels for all samples in the dataset, we reran rMATS for all samples with the command-line option “–cstat 0”.

The list of RNA binding proteins was obtained from the rMAPS website (http://rmaps.cecsresearch.org/Help/RNABindingProtein). In addition to the RNA binding motifs that were tested using the default settings on the rMAPS website, we also conducted tests on additional UGG-enriched motifs that have been previously identified as binding sites for the RNA binding proteins ESRP1 [73,74] and ESRP2 [75] (Table S6). Following Yang et al. [64] and the CISBP-RNA database [76] (http://cisbp-rna.ccbr.utoronto.ca), we assumed that the proteins RBFOX1 and RBFOX2 both bind to the same mRNA motif ([AT]GCATG[AC]).

## ACKNOWLEDGMENTS

We wish to thank Yochai Israelashvili, Naomi Pode-Shakked, Iddo Ben-Dov, Thomas Tuschl, and all members of our lab for helpful comments and suggestions. The results published here are in whole or part based upon data generated by the Therapeutically Applicable Research to Generate Effective Treatments (TARGET) initiative, phs000218, managed by the NCI. The data used for this analysis are available through the GDC (https://gdc.cancer.gov/about-data/publications#/?groups=&years=&programs=TARGET&order=desc). Information about TARGET can be found at https://www.cancer.gov/ccg/research/genome-sequencing/target/about.

## AUTHOR CONTRIBUTIONS

Study initiation and conception – Y.T. and T.K.; Data analysis – Y.T. and T.K.; Other intellectual contribution – A.U., B.D., P.H., J.G.; Manuscript writing – Y.T and T.K.

**Supplementary information:** Supplementary text and figures.

**Table S1:** An excel file containing: (i) raw gene expression counts (ii) normalized and modified log-transformed gene expression counts (that is, log2(1+x) where x represents the DESeq2-normalized counts), where genes with zero expression in all samples were omitted (iii) gene expression profiles of the three archetypes (iv) a table containing merged metadata from the TARGET database.

**Table S2:** Gene Ontology (GO) enrichment analysis for the stromal, blastemal, and normal kidney archetypes. For each archetype, genes for which log2FC > 3 with respect to the other two archetypes were selected and inserted to ToppGene.

**Table S3:** A list of 102 selected genes that are known from the literature to be associated with kidney development and tumorigenesis.

**Table S4:** An excel file containing: (i) Lists of exons that were found by rMATS to be significantly alternatively spliced (FDR = 0, |Δ*ψ*| > 0.1) in the three comparisons between samples located near the three archetypes (ii) Lists enriched gene ontologies found by ToppGene for each comparison.

**Table S5:** A table of inclusion levels for transcripts that were found to be significantly alternatively spliced (FDR = 0, |Δ*ψ*| > 0.2) in the three comparisons between samples located near the three archetypes. This table includes the inclusion levels for all samples in the dataset. Only SE (single exon) and MXE (mutually exclusive exons) splicing events were included since these were the majority.

**Table S6:** A list of known RNA binding proteins and motifs that were examined in this study. **Program:** A compressed directory containing programs and datasets for reproducing the main results.

